# A brain atlas of the camouflaging dwarf cuttlefish, *Sepia bandensis*

**DOI:** 10.1101/2022.01.23.477393

**Authors:** Tessa G. Montague, Isabelle J. Rieth, Sabrina Gjerswold-Selleck, Daniella Garcia-Rosales, Sukanya Aneja, Dana Elkis, Nanyan Zhu, Sabrina Kentis, Frederick A. Rubino, Adriana Nemes, Katherine Wang, Luke A. Hammond, Roselis Emiliano, Rebecca A. Ober, Jia Guo, Richard Axel

## Abstract

The coleoid cephalopods (cuttlefish, octopus, and squid) are a group of soft-bodied marine mollusks that exhibit an array of interesting biological phenomena, including dynamic camouflage, complex social behaviors, prehensile regenerating arms, and large brains capable of learning, memory, and problem-solving [1–10]. The dwarf cuttlefish, *Sepia bandensis*, is a promising model cephalopod species due to its small size, substantial egg production, short generation time, and dynamic social and camouflage behaviors [11]. Cuttlefish dynamically camouflage to their surroundings by changing the color, pattern and texture of their skin. Camouflage is optically-driven, and is achieved by expanding and contracting hundreds of thousands of pigment-filled saccules (chromatophores) in the skin, which are controlled by motor neurons emanating from the brain. We generated a dwarf cuttlefish brain atlas using magnetic resonance imaging (MRI), deep learning, and histology, and we built an interactive web tool (cuttlebase.org) to host the data. Guided by observations in other cephalopods [12–20], we identified 32 brain lobes, including two large optic lobes (75% the total volume of the brain), chromatophore lobes whose motor neurons directly innervate the chromatophores of the color-changing skin, and a vertical lobe that has been implicated in learning and memory. This brain atlas provides a valuable tool for exploring the neural basis of cuttlefish behavior.

## Results

### An MRI-based 3D brain atlas of the dwarf cuttlefish

The dwarf cuttlefish, *Sepia bandensis*, is a small, tropical species from the Indo-Pacific (Figure 1A) that exhibits a rich repertoire of behaviors. The skin of the dwarf cuttlefish is covered in hundreds of thousands of chromatophores (pigment-filled saccules surrounded by radial muscles) whose expansion states are controlled by motor neurons projecting from the brain [21]. During camouflage (Figure 1B), conspecific communication (Figures 1C,D), and deimatic behavior, dwarf cuttlefish dynamically alter the color, pattern and texture of their skin using chromatophores and subcutaneous papillae. This process is driven by visual circuits in the brain, and is therefore extremely rapid, occurring in milliseconds [22, 23]. Thus, the skin patterning of cuttlefish reflects both the organism’s perception of the external world and its internal state.

**Figure 1.**
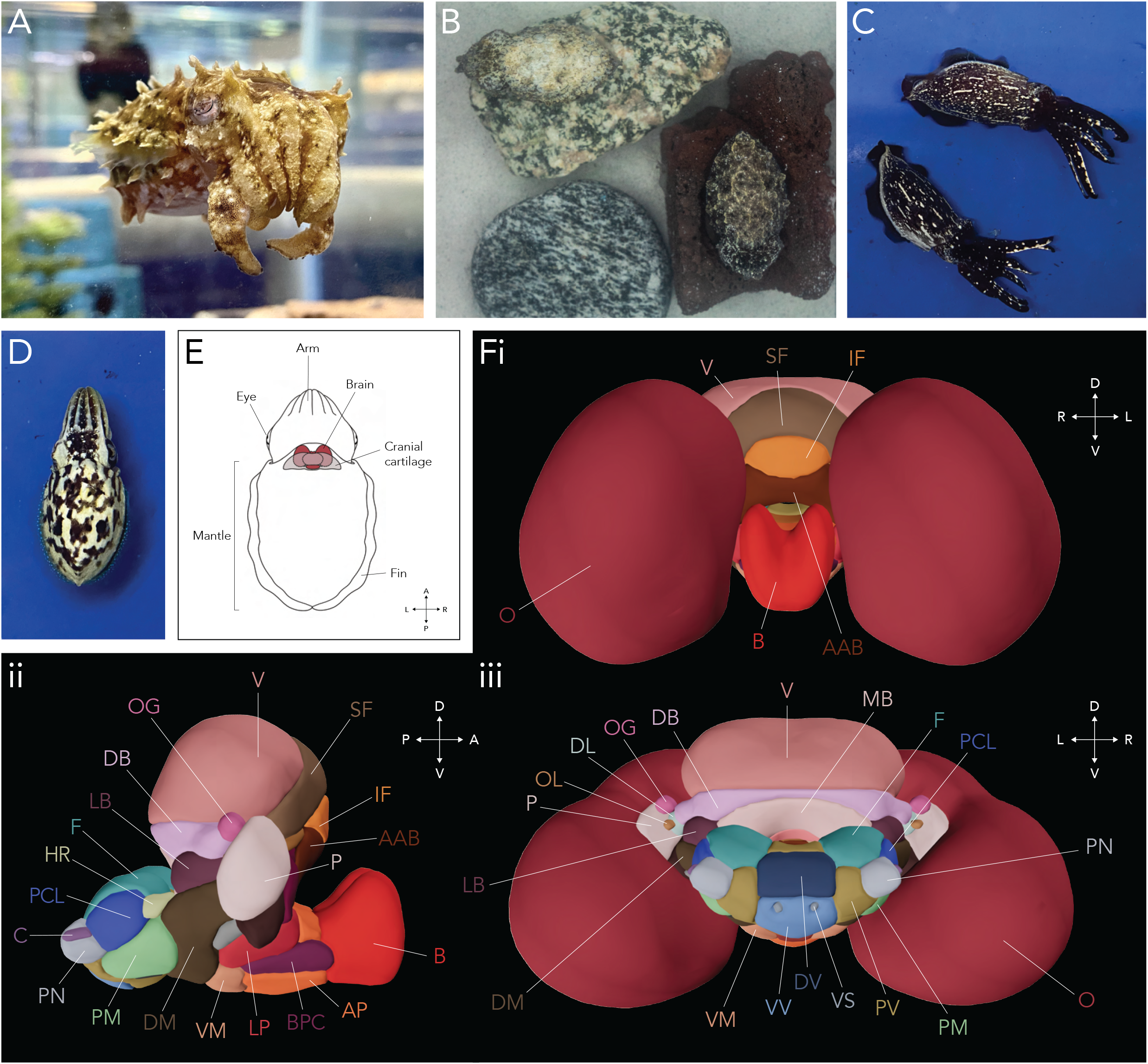
3D brain atlas of the dwarf cuttlefish, *Sepia bandensis*. A) Adult dwarf cuttlefish (~8 cm in length). B) Cuttle-fish camouflage to their surroundings by changing the color and texture of their skin. C) The innate skin pattern and posture adopted by male dwarf cuttlefish during aggressive interactions. D) An innate skin pattern frequently adopted by dwarf cuttlefish in the presence of conspecifics. E) Anatomy of a cuttlefish. The brain is encased on its posterior side by rigid cartilage. F) 3D template brain, based on MR imaging of 8 brains (4 males, 4 females) and manual annotations of 32 brain lobes and 12 nerve tracts. For abbreviations, see Table 1. (i) Anterior view, (ii) Right view (shown without the optic lobes) and (iii) Posterior view.

The dwarf cuttlefish brain is located posterior and medial to the eyes and is encased on the posterior side by cranial cartilage (Figure 1E). We performed *ex vivo* magnetic resonance imaging (MRI) of the brains of 8 adult dwarf cuttlefish (4 males, 4 females) at 50 μm isotropic resolution and then combined manual segmentation with deep learning techniques to extract each brain from its surrounding tissue (see Methods, Figure S1). Using 6 independent annotators, we segmented the brain lobes using prior neuroanatomical descriptions as a guide [12, 18, 19]. Finally, we co-registered the 8 brains to create a merged, annotated template brain (Figure 1F).

The *ex vivo* dwarf cuttlefish brain is 94% the volume of the *ex vivo* mouse brain (Table S1, Methods), which is consistent with the cuttlefish possessing a deep behavioral repertoire, akin to that of a small mammal. The cuttlefish brain surrounds the esophagus, and can been coarsely divided into a supraesophageal mass and a subesophageal mass [24, 25] that we further divided into 32 discrete lobes (Table 1). This value is in accord with previous anatomical studies in other species, but it remains possible that the annotated lobes can be subdivided further. The largest brain lobes, the optic lobes (O), receive direct projections from the retina and are engaged in visual processing [13] (Figures 1Fi,iii). They comprise 75% of the total volume of the brain (Table S1), consistent with the dominant role of vision in guiding many cephalopod behaviors [26]. One relevant projection of the optic lobe is the lateral basal lobe (LB), an intermediate station in the camouflage pathway [12, 13, 18, 20]. The lateral basal lobe may use processed visual information from the optic lobes to compute the most appropriate skin pattern components needed for camouflage. The vertical lobe complex lies on the dorsal side of the supraesophageal mass (Figure 1F). This complex is considered to be the learning and memory center of the brain [27] and may facilitate short-term [28], spatial [29], and visual [30] learning in cephalopods, including working [31] and episodic-like memory [32]. The vertical lobe (V) is directly underneath the dorsal cranial cartilage – an ideal structure for anchoring an electrode or GRIN lens for neural recordings (Figure 3B).

**Table 1.**
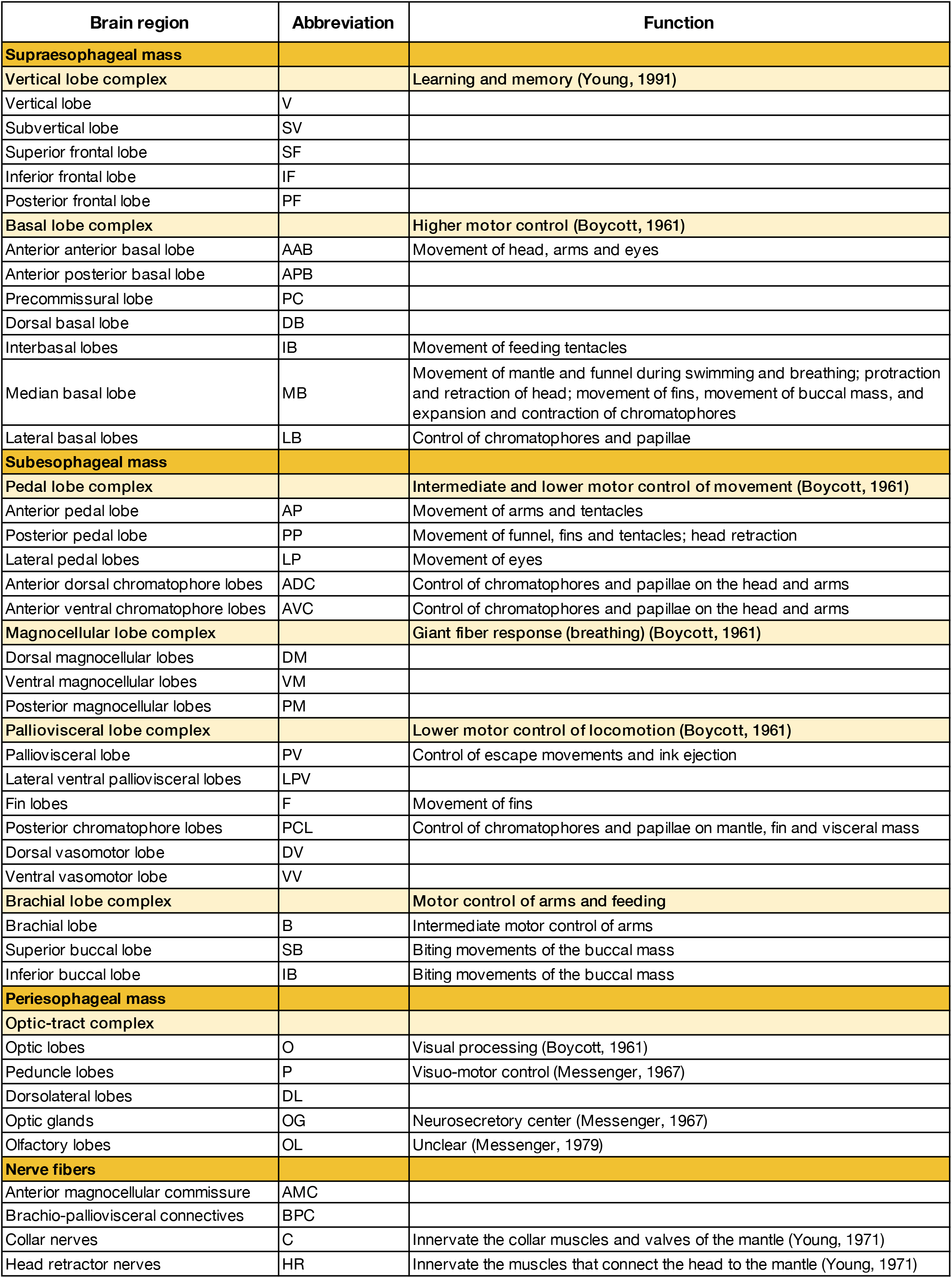
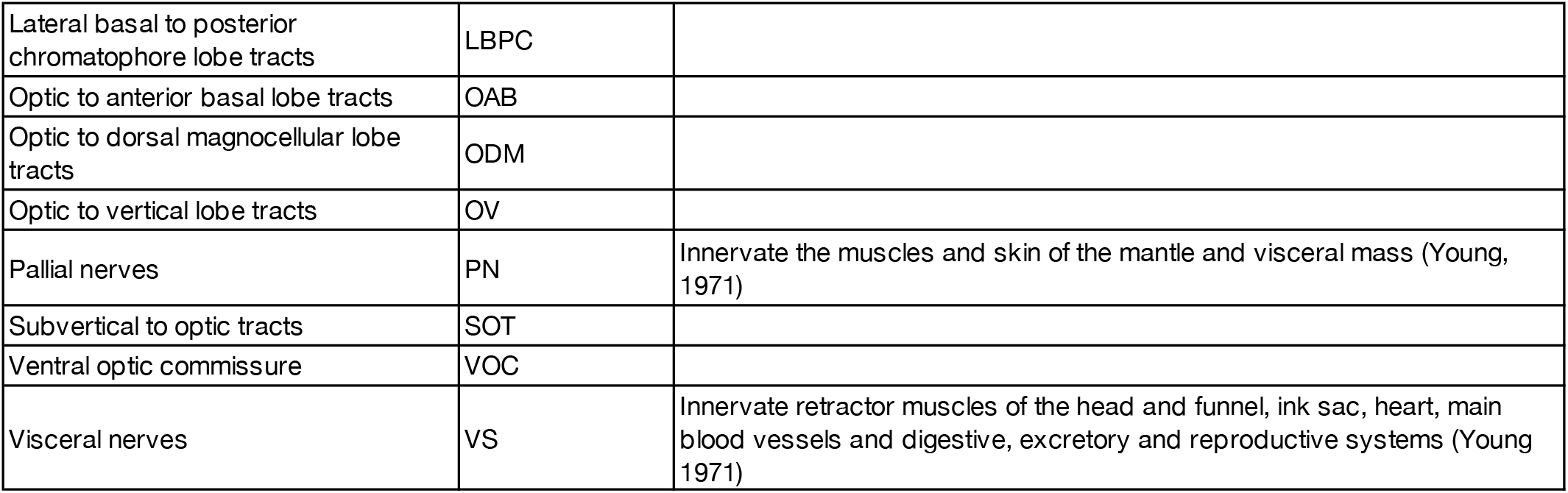
The lobes of the cuttlefish brain.

The subesophageal mass comprises multiple structures that elicit simple motor actions. There is a rough mapping of lobe position in the brain to motor target in the body. For instance, the lobes in the anterior portion of the subesophageal mass control movements of structures in the anterior portion of the cuttlefish: the brachial lobe (B) controls the arms, the lateral pedal lobes (LP) control movements of the eyes (Figure 1Fii), and the anterior chromatophore lobes control the chromatophores of the head and arms [18]. Connections between the anterior and posterior chromatophore lobes are thought to coordinate skin patterns between the head, arms and body.

The posterior region of the subesophageal mass (the posterior subesophageal mass, PSM), projects through an opening in the cranial cartilage (Figure 3B). Structures in this region control motor actions in the more posterior aspect of the cuttlefish. For instance, the posterior chromatophore lobes (PCL) control the chromatophores of the mantle, the fin lobes (F) control movement of the fins, and the palliovisceral lobe (PV) controls escape movements and inking, which are mediated by the funnel [18] (Figure 1Fiii). Some cephalopod brain lobes have been assigned function through lesion [29, 33–37] and electrophysiological studies [18, 38–43]. However, the functions of most brain regions are unknown.

### A histological brain atlas for the dwarf cuttlefish

We complemented the anatomical description of the cuttlefish brain with histological examination of the entire brain to obtain cellular resolution. We sectioned the brain in the transverse, horizontal and sagittal planes (Figure 2A) and stained the sections with Phalloidin, an F-actin peptide that labels axons, and NeuroTrace, a Nissl stain that labels neuronal cell bodies and glia. Our annotated 3D MRI datasets and prior neuroanatomical descriptions [12, 18, 19] were used to describe the histological organization of 32 brain lobes and 12 nerve tracts (Figures 2B-D).

**Figure 2.**
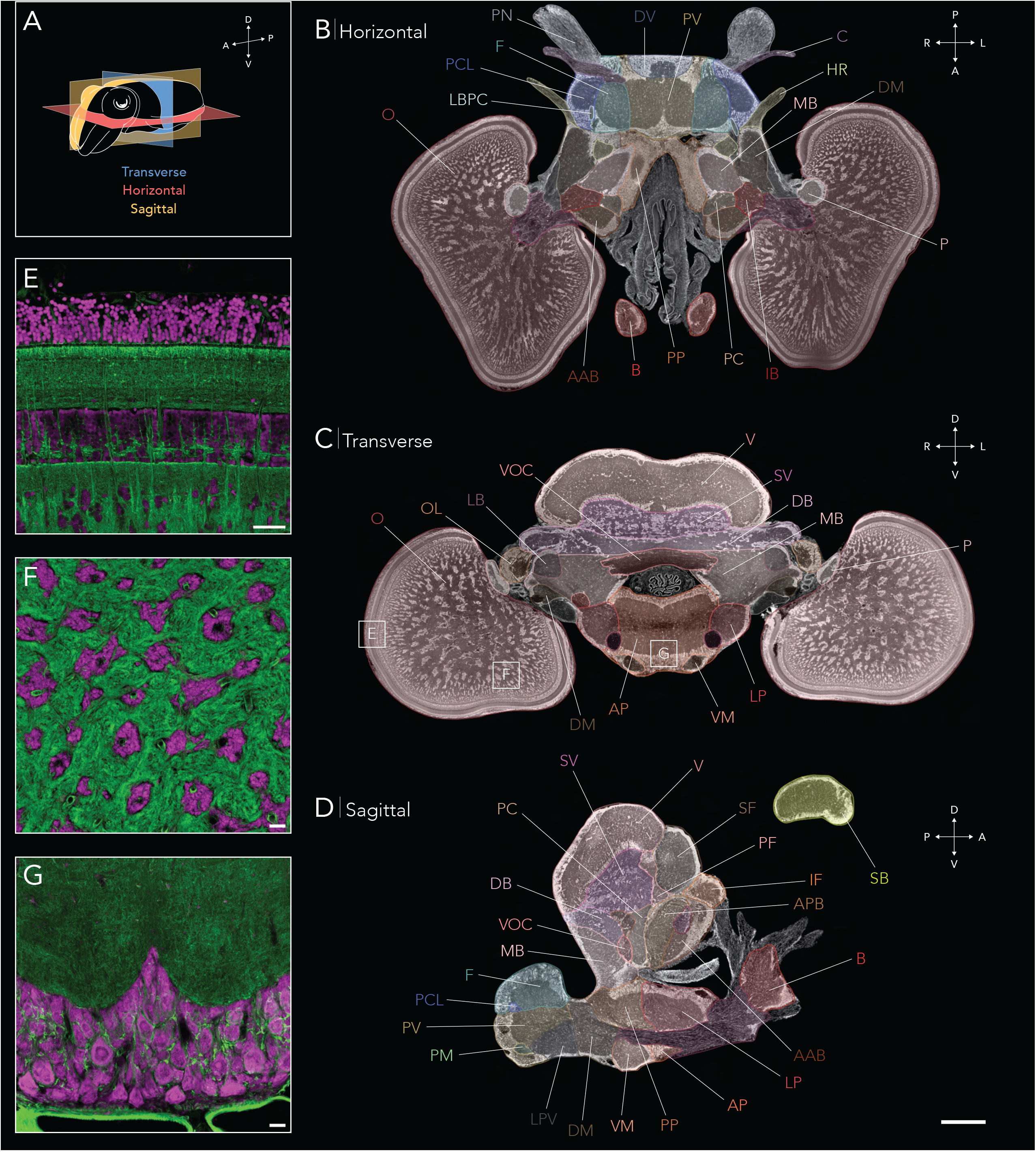
Histological brain atlas of the dwarf cuttlefish, *Sepia bandensis.* A) The dwarf cuttlefish brain was sectioned in the transverse, horizontal and sagittal planes. B-D) Representative histological slices of the cuttlefish brain stained with NeuroTrace, a Nissl stain that labels cell bodies, and annotated with 32 brain lobes and 12 nerve tracts. Scale bar, 1 mm. For abbreviations see Table 1. E-G) Confocal images of cuttlefish tissue stained with NeuroTrace (purple) and Phalloidin (an F-actin peptide that labels axons, green). Boxed areas in C show approximate locations of confocal images within the brain. E) Optic lobe layered cortex, F) Optic lobe medulla, G) Anterior pedal lobe. Confocal scale bars, 50 μm.

Phalloidin and NeuroTrace staining confirmed previously described organizational features of the cephalopod brain. The majority of lobes are discrete and bounded, a feature not uniformly observed in vertebrate brains: each lobe contains an outer perikaryal layer of cell bodies that surrounds a dense, inner neuropil (Figures 2B-D). Dwarf cuttlefish neuronal cell bodies range in diameter from ~5 um (e.g. the vertical and optic lobes) to <80 um in some motor areas (e.g. the fin and pedal lobes) (Figures 2E-G).

Similar to other coleoid cephalopods, the *Sepia bandensis* optic lobe consists of two ordered outer layers, the inner and outer granular layers, apposing a zone of fibers (Figure 2E) [13, 20, 44]. Because the cephalopod retina contains only photoreceptors, the outer layers of the optic lobe may function like the vertebrate retina and perform the initial stages of visual processing [44, 45]. Beneath the outer layers, the cephalopod optic lobe features a novel neural organization: a central medulla that features cell ‘islands’ (Figure 2F) connected in an elaborate tree-like structure [13, 20, 46]. Cell bodies near the cortex (the ‘branches’) appear to form columns (Figures 2B,C), which may reflect the presumed retinotopic order of the outer layers [13, 20, 47].

Five pairs of nerves project from the cuttlefish brain that are easily distinguishable in histological slices and contain many of the brain’s major efferents. The largest nerves, the pallial nerves, exit the posterior side of the brain and contain motor fibers that innervate the chromatophores of the mantle skin and the muscles of the mantle and fin (Figure 2B) [12]. The motor neurons that originate in the posterior chromatophore lobe exit via the pallial nerve, pass through the stellate ganglion, and directly innervate the radial muscles of the chromatophores [48]. Severing the pallial nerve results in blanched skin, loss of the skin’s textural control, and a limp fin on the ipsilateral side of the body [49]. Closely apposed to the pallial nerves are the collar nerves, which innervate the collar muscles and valves of the mantle (Figure 2B) [12]. The visceral nerves are positioned near the midline on the posterior side of the brain and innervate the heart, ink sac muscles, digestive, reproductive and excretory systems [12]. Finally, the head retractor nerves, positioned ventral to the lateral basal lobes, innervate the muscles that connect the head to the mantle (Figure 2B) [12]. When dwarf cuttlefish sense danger, they can retract their head inside their body using these muscles.

### Cuttlebase - a web tool for visualizing the cuttlefish brain

We built an interactive web tool, Cuttlebase, to maximize the utility of our brain atlas (Figure 3A). Cuttlebase features multiple tools for the dwarf cuttlefish, including the annotated histological sections of the brain in 3 planes (Figure 3A), the 3D brain model described above (Figure 3B), and a 3D model of an entire adult cuttlefish labeled with 26 organs, including the animal’s three hearts, gills, ink sac, beak, cuttlebone and digestive system (Figure 3C). Furthermore, the 3D body model features the eight brachial nerves that innervate the cuttlefish’s arms, and the two pallial nerves, which exit the posterior aspect of the brain, circumscribe the digestive gland, and then pass through a hole in the mantle musculature to reach the stellate ganglion [50]. Cuttlebase is easy-to-use, with an array of features including responsive color-coded labels for each brain region, a dynamic scale bar, the ability to zoom, rotate and screenshot the data, and synchronized graphics that denote the brain’s orientation.

**Figure 3.**
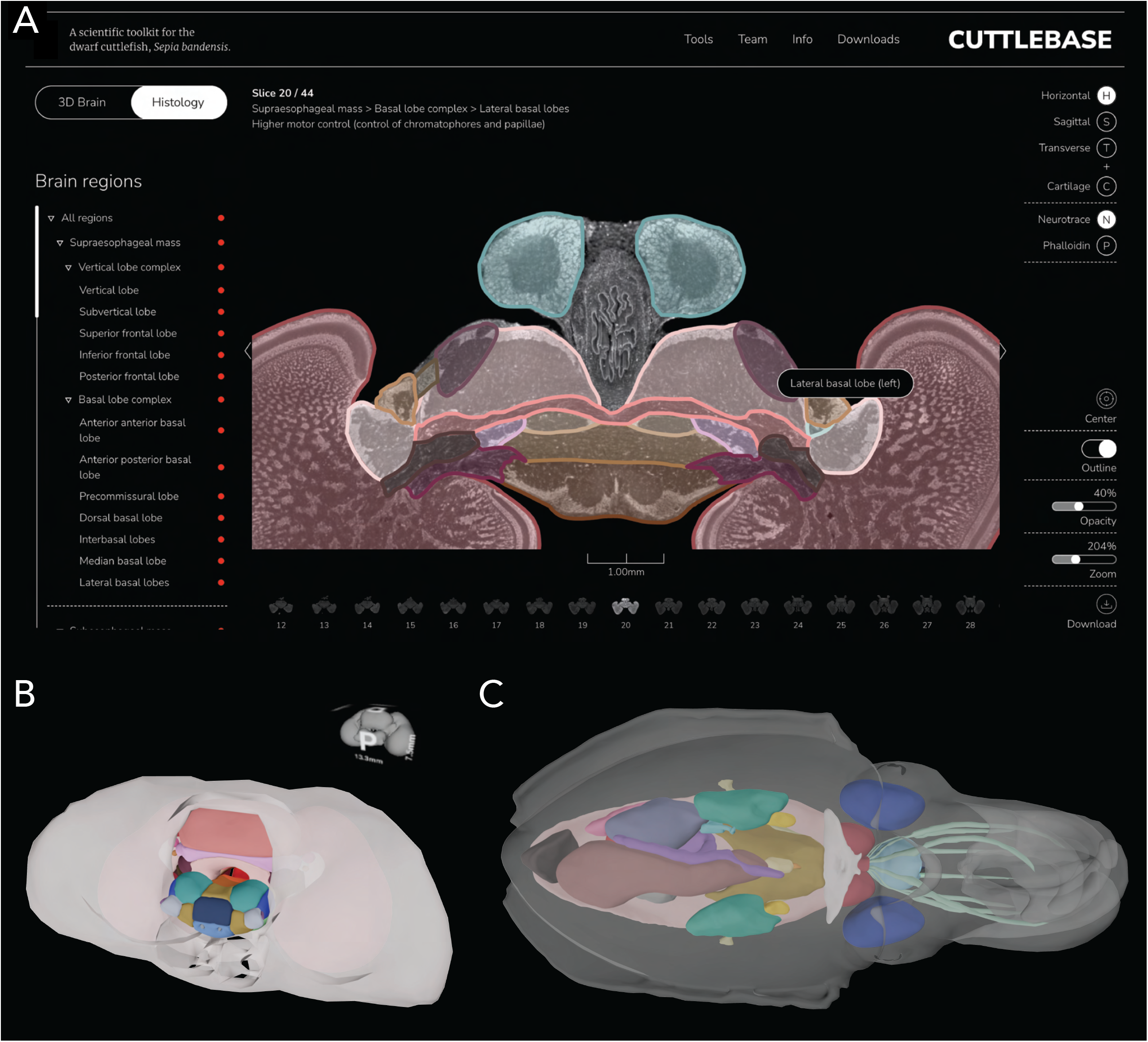
Cuttlebase is a scientific toolkit for the dwarf cuttlefish. A) Cuttlebase hosts multiple dwarf cuttlefish tools including an interactive histological brain atlas, which features color-coded and responsive labels, a dynamic scale bar, and the ability to screenshot, zoom and rotate the data. B) The 3D brain atlas can be visualized with the cranial cartilage and features a synchronized graphic that denotes the brain’s orientation. C) Cuttlebase also features a body atlas, labeled with 26 organs and nerves.

Cuttlebase was designed for both educational and research settings. For its use in education, we added features that increase accessibility, including: (i) A landing page that introduces cuttlefish behavior, brain anatomy, evolution and animal care; (ii) An info page that shows the cuttlefish’s gross anatomy and the sectioning planes used in the histological atlas; (iii) An option to show the 3D brain model in the context of the cuttlefish’s body; and (iv) Mouseover descriptions of the function of each brain lobe. For expert users, we included an array of more advanced features. First, we included a dynamic scale bar in the histology mode to help users measure distances within the brain and devise brain coordinates for stereotactic surgeries. Second, we imaged the cranial cartilage — the only rigid tissue around the brain — and provided its interactive graphic (Figure 3B) to facilitate the development of methods to head-fix cuttlefish, which is an important step for stabilizing the brain for two-photon imaging or electrophysiology. Finally, we made the MRI data, brain model and full-resolution histology data available to download, so users can carry out their own MRI analyses, annotate the histological images with the results of tracing experiments, and import the to-scale 3D cuttlefish into modeling and 3D-printing software.

## Discussion

The 3D and histological brain atlases presented here serve multiple functions. First, they provide a comprehensive resource for comparative neuroanatomical analyses. Little is currently known about the biology of the dwarf cuttlefish (*Sepia bandensis*), whereas the common cuttlefish (*Sepia officinalis*) has been studied extensively to understand behavior [8], learning and memory [1] and the neural control of skin patterning [4, 23]. Comparison of our dataset with the neuroanatomy of the common cuttlefish [18, 51] and mourning cuttlefish, *Sepia plangon* [52], reveals substantial neuroanatomic resemblance despite the species’ different sizes, behaviors and habitats [11, 53]. Every cuttlefish species has evolved a different ensemble of skin pattern components for camouflage and social communication. *Sepia bandensis*, which live in Indo-Pacific coral reefs, assemble skin patterns with predominantly high frequency patterns, likely reflecting the visual statistics of their tropical habitat. By contrast, *Sepia officinalis*, which live in the rocky waters of Europe, include in their repertoire a set of large, disruptive patterns that resemble rocks [53]. During male social encounters, although *Sepia officinalis* and *Sepia bandensis* both exhibit a high-contrast, stereotyped aggression pattern, their precise manifestations differ: *Sepia officinalis* create a zebra pattern [53], whereas the *Sepia bandensis* aggression pattern is stippled (Figure 1B) [11]. Additionally, although both species are capable of creating dynamic waves across their skin, they do so in opposite directions (Video 1) [10]. Such fine-scale differences as these are unlikely to be reflected at the coarse anatomical level. Spatial transcriptomics, fine anatomical tracing and developmental tracking may reveal how the neural circuits diverged to facilitate different skin patterns.

Furthermore, the investigation of the neural circuits underlying camouflage may reveal how the optic lobe-lateral basal lobe-chromatophore lobe pathway creates an internal representation of a visual scene, and then recreates an approximation of the scene on the skin. Recording neural activity in the camouflage circuit, as well as precise anterograde and retrograde tracing studies, may answer questions such as: *Is the cuttlefish capable of only a limited number of patterns that when combined recapitulate the natural scene? Is the neural network underlying camouflage capable of exhibiting experience-dependent plasticity? Are the activity patterns in the chromatophore lobe topographic with patterns on the skin, or are they an abstraction? How is an internal, abstract representation of the visual world transformed to a more figurative representation on the skin?*

Comparison of the dwarf cuttlefish brain with other decapodiformes (10-limbed cuttlefish and squid) and octopodiformes (8-limbed octopus and vampire squid) reveals substantial organizational similarities and a few key differences. First, unlike the octopodiformes, the decapodiformes possess a giant-fiber system, which is used for funnel-based jet propulsion during escape behaviors. The giant cells are located in the ventral magnocellular lobe, and receive input from the optic lobes and statocysts (the cephalopod vestibular system) [54, 55]. Second, octopodiformes employ sophisticated use of their arms and demonstrate tactile learning; their brachial, pedal and inferior frontal lobes are larger than the decapodiformes [56]. Third, the octopus vertical lobe is folded into gyri, creating a larger surface area [12], whereas the cuttlefish vertical lobe is dome-shaped and lacks gyri. Finally, the decapodiformes, but not octopodiformes, possess a pair of fin lobes used for fin locomotion [18]. Most decapodiformes use two forms of locomotion: jet propulsion using the funnel, and swimming using the funnel and fins [57]. Interestingly, the dwarf cuttlefish uses an additional mode: it can walk bipedally using its ventral arms when combined with water ejection through the funnel (Video 2). The gross morphology of the dwarf cuttlefish brachial and pedal lobes are not readily distinguishable from other cuttlefish species, but deeper analysis of the circuitry might uncover the evolution of a control system for coordinated locomotion.

This atlas also provides a valuable resource for comparisons across phyla. For instance, despite ~600 million years of separation between cuttlefish and fruit flies [58], the brains of these species share coarse organizational features. The esophagus runs through the center of the brain, two large optic lobes flank the central brain mass, and the learning and memory centers - the mushroom body in *Drosophila* and the vertical lobe in cuttlefish - lie on the dorsal side of the brain. Furthermore, the two outer layers of the cuttlefish optic lobe may share properties with the lamina and medulla of the dipteran visual system [59]. Interestingly, recent single-cell gene expression profiling of a developing octopus brain has revealed molecular resemblances between the Kenyon cells of the *Drosophila* mushroom body and the vertical lobe cells of the octopus brain [60].

The dwarf cuttlefish brain atlas presented here serves as both an educational resource and a tool for the investigation of the neural basis of cephalopod behavior. Users can employ Cuttlebase to plan the target angle and depth of electrode insertion or to design head plates that anchor GRIN lenses for calcium imaging. Following the completion of *in vivo* experiments, the histological atlas affords users the ability to identify the experimentally targeted brain regions by directly comparing post-operative tissue slices with the atlas. Beyond the interrogation of neural circuits by imaging, recording or tracing, the histological atlas can be used to assign function to brain lobes by mapping immediate early gene expression following a behavioral paradigm, and to map spatially-restricted gene expression to neuroanatomy. The recent generation of squid and octopus cell atlases by single-cell RNA-seq complement the generation of neuroanatomical atlases, providing a more comprehensive view of brain organization [60–63].

Finally, it is important to note that this atlas represents a first attempt to demarcate the brain lobes of the dwarf cuttlefish, but these demarcations will likely need to be adjusted as we learn more about the functional organization of the cephalopod brain. At present, the vast majority of functions assigned to the cuttlefish brain lobes originated from a single study, in which brain lobes were systematically stimulated by hand [18]. The application of new technologies, such as transgenic tools, will likely lead to a more nuanced view of lobe function, and the development and application of retrograde and anterograde tracing methods will improve our understanding of connectivity within the brain.

In summary, Cuttlebase was designed to aid the investigation of cuttlefish biology, as well as foster the curiosity of scientists and non-scientists alike. The generation of the neuroanatomical and histological atlases for the dwarf cuttlefish may inform studies in other cephalopods, for which atlases do not exist, and now renders *Sepia bandensis* a more facile system for the study of the neural control of camouflage.

## Acknowledgements

We wish to thank Connor Gibbons, Sonia Thomas, Telicia Lewis, Sarah Wilson and Josh Barber for cuttlefish care, the Zuckerman Institute Cellular Imaging platform for instrument use and technical support, Lokke Highstein for IT support, and Gwenyth Card and Carrie Albertin for valuable discussions. This project was funded by a Zuckerman Institute MRI Seed Grant (T.G.M. and R.A.), the Howard Hughes Medical Institute Hanna H. Gray Fellowship (T.G.M.), the Zuckerman Institute BRAINYAC Program (R.E.) and the Howard Hughes Medical Institute (R.A.)

## Author contributions

T.G.M., R.A.O. and J.G. conceived the project. T.G.M., I.J.R. and J.G. performed MRI experiments. T.G.M. performed histology experiments. S.G-R. conducted MRI data processing steps, with input from N.Z. and J.G. The MRI data was annotated by I.G.R., T.G.M., D.G-R., S.K., F.A.R., A.N., K.W., and R.E. The histology data was annotated by I.J.R. and T.G.M. L.A.H. developed the BrainJ histology data processing software. D.E. designed and S.A. developed the Cuttlebase website, respectively. J.G. and R.A. provided supervision. T.G.M. wrote the original draft of the manuscript. T.G.M. and R.A. reviewed and edited the manuscript. T.G.M. and R.A. acquired funding for the project.

## Methods

### Ethics statement

The use of cephalopods in laboratory research is currently not regulated in the USA. However, Columbia University has established strict policies for the ethical use of cephalopods, including operational oversight from the Institutional Animal Care and Use Committee (IACUC). All of the cuttlefish used in this study were handled according to approved IACUC protocols (AC-AABE3564), including the use of deep anesthesia and the minimization and prevention of suffering.

### 3D brain atlas

#### MRI acquisition

8 adult cuttlefish (4 male, 4 female) were euthanized in 10% ethanol, and the brain and connected eyes were surgically removed and fixed overnight in a solution of 4% paraformaldehyde (PFA) in filtered artificial seawater (FASW) at 4°C. The fixed brains were washed in PBS, incubated in 0.2% OMNISCAN (gadodiamide) for 2 days at 4°C to increase contrast, and then suspended in fomblin in a 15 mL conical tube. Imaging was performed on a Bruker BioSpec 94/30 horizontal small animal MRI scanner (field strength, 9.4 T; bore size, 30 cm) equipped with a CryoProbe and ParaVision 6.0.1 software (Bruker). A 23 mm 1H circularly polarized transmit/receive-capable mouse head volume coil was used for imaging. For each cuttlefish, one scan was acquired of the brain and eyes (to obtain a scan of the entire cranial cartilage), and a higher resolution scan was acquired of the brain only. T1-weighted images were acquired with a Fast Low Angle Shot (FLASH) sequence (brain/eye scan: TR = 50 ms, TE = 10 ms, FOV = 30 x 24 x 15 mm^3^, voxel size = 100 x 100 x 100 μm^3^, scan time = 33 m 12 s; brain-only scan: TR = 50 ms, TE = 8.5 ms, FOV = 14 x 12 x 11 mm^3^, voxel size = 60 x 60 x 60 μm^3^, scan time = 4 hr 28 m).

#### MRI processing & brain extraction

All scans underwent N4 bias field correction [64]. Whole brain scans were isotropically upsampled to 50 μm isotropic resolution with cubic B-spline interpolation. To computationally extract the brains from their surrounding tissue, brain masks were generated by an in-house deep learning model [65], which was pre-trained with brain masks manually annotated in 3D Slicer [66]. The deep learning brain masks were manually polished using the 3D Slicer Segment Editor and then used to extract the brain from each brain-only MRI scan (“brain-extracted images”) (Figure S1). For the brain/eye scans, the cranial cartilage was manually segmented in 3D Slicer using the Segment Editor. The cartilage masks were used to extract the cartilage from each brain/eye MRI scan (“cartilage-extracted images”).

#### Segmentation

Two of the whole brain scans (1 male, 1 female) were manually segmented in 3D Slicer by 6 independent annotators guided by prior neuroanatomical descriptions [18, 19]. The brain label maps for each subject were merged using pixel-level majority voting, transformed to the remaining 6 subjects, and then manually corrected, resulting in 8 brain label maps corresponding to the 8 subjects. Lobe locations were determined by prior annotations [18, 19] and precise lobe boundaries were estimated using the location of an outer cell body layer, which is a feature of most cephalopod brain lobes. The combined use of high-resolution histological data and three-dimensional MRI data aided the discrimination of lobes in 3D space.

#### Generation of the template brain

The final MRI atlas was built by merging the brain template, built from the high-resolution brain-only scans and their mirror images, with the cranial cartilage template, built from the brain/eye scans. First, to generate each template, a population average of brain-extracted or cartilage-extracted images was constructed through an iterative process by averaging the co-registered images over multiple cycles using a symmetric diffeomorphic registration algorithm [67]. The brain label map (annotations) for each subject was diffeomorphically transformed to the whole brain template space and combined through pixel-level majority voting. The brain/eye template was isotropically upsampled to match the 50 μm resolution of the whole brain scans, and the whole brain template underwent rigid registration to align it with the brain/eye template. Finally, the brain regions of the whole brain template were combined with the cartilage regions of the whole head template to build a single atlas. In the regions where the two templates overlapped, pixel values of the high-resolution whole brain template were selected. The cuttlefish brain label map was smoothed by taking a majority vote in a local neighborhood with a 3 x 3 x 3 kernel size.

#### Brain volume calculation

A mouse MRI brain model (15 μm resolution, average of 18 *ex vivo* subjects) was downloaded from the Australian Mouse Brain Mapping Consortium (nonsymmetric version, NiFTI format, https://imaging.org.au/AMBMC/Model), and then downsampled to 50 μm resolution and imported into 3D Slicer. Using the Editor tool at a threshold of zero, a mask was generated of the entire mouse brain, and the spinal cord was manually removed using the eraser tool. The volume of the mouse brain mask was calculated using the Label Statistics tool (total volume: 332.7 mm^3^). To calculate the volume of the cuttlefish brain, the volume of each brain lobe in the merged, template brain was calculated using the Label Statistics tool and summed (total volume: 312.1 mm^3^), see Table S1.

### Histological atlas

Adult cuttlefish were euthanized in 10% ethanol, and the brain and connected eyes were surgically removed and fixed overnight in a solution of 4% paraformaldehyde (PFA) in filtered artificial seawater (FASW) at 4°C. Note that fixing brains in a solution of PBS (instead of FASW) created abnormal cell morphologies. The fixed brains were washed in PBS, the eyes were removed with a scalpel, and the brains were incubated in 10% sucrose overnight followed by 30% sucrose overnight at 4°C. The brains were embedded in OCT on dry ice and stored at −80°C. Each brain was sliced in 100 μm sections on a cryostat (Leica CM3050 S), and the sections were dried overnight at room temperature. The sections were stained with Phalloidin (Life Technologies #A12379, 1/40 dilution) and NeuroTrace (Life Technologies #N21482, 1/20 dilution) and then imaged on a custom-built Nikon AZ100 Multizoom Slide Scanner. Images were registered using BrainJ (http://github.com/lahammond/BrainJ). Brightness and contrast were adjusted uniformly across each image, and surrounding tissue was removed manually from the image in FIJI and Photoshop. The data was manually segmented in 3D Slicer by two annotators using a neuroanatomical study [18] and our 3D brain data for reference.

### Whole body atlas

An adult male cuttlefish was anesthetized in MgCl2 (17.5g/L) and then euthanized in 10% ethanol, fixed for 2 days in 4% PFA/FASW at 4°C, and then transferred to 0.2% OMNISCAN (gadodiamide) for 2 days at 4°C to increase contrast. The fixed specimen was suspended in fomblin in a custom-made vessel and imaged on a Bruker BioSpec 94/30 horizontal small animal MRI scanner (field strength, 9.4 T; bore size, 30 cm) equipped with a CryoProbe and ParaVision 6.0.1 software (Bruker). A 112/86-mm 1H circularly polarized transmit/receive-capable volume coil was used for imaging. T1-weighted images were acquired with a FLASH sequence (TR = 55 ms, TE = 17 ms, FOV = 90 x 48 x 38 mm^3^, voxel size = 100 x 100 x 100 μm^3^, scan time = 2 hr 47 m). The scan underwent N4 bias field correction [64] and was manually segmented in 3D Slicer using anatomical descriptions [68].

### Cuttlebase

The Cuttlebase web content is delivered using React, a Javascript front-end framework for dynamic websites. React-three-fiber (a Three.js wrapper for React) is used to assist with interactions in the 3D view.

#### 3D brain

In 3D Slicer, each segment (brain lobe or tract) of the final, template brain was exported as an STL file and then labelled with a unique identifier in Blender, a 3D authoring software. The 3D model was exported as a GLB, and then imported into a webpage using Three.js - a Javascript library for handling 3D content on the web. A custom web-interface was created to assign colors to each of the region meshes, and this data was exported as a JSON file.

#### Histology

To convert the histological annotations to high-resolution images that could be toggled on Cuttlebase, each segment (brain lobe or tract) for each brain (horizontal, sagittal and transverse) was converted to a binary label map in 3D Slicer and saved as a TIFF stack. Each TIFF stack was resized to the canvas size of the original image in Fiji [69], and saved as a JPEG Stack in monochrome. The images were then inverted and processed with Potrace (through a custom Node.js script), to create SVGs for each region of each layer. The SVGs for all regions in a single layer were combined, and each region was assigned the color corresponding to the 3D atlas. These images, along with the originals, were resized and cropped for more efficient web delivery. Additionally, the cartilage in each Phalloidin section was isolated (by outlining) using Adobe Illustrator, and exported as a PNG. These images were used as masks on the NeuroTrace layers with the command-line tool ImageMagick, to automate this process for the remaining images. Custom Bash scripts were used to manage and organize the large amounts of data and the processing steps required.

#### Data availability

All data generated in this study is available to download from cuttlebase.org/downloads.

## Video legends

**Video 1**. Dwarf cuttlefish produce waves (of unknown function) on their skin.

**Video 2**. Dwarf cuttlefish can walk bipedally using their ventral arms when coupled with water ejection through the funnel.

---

**Figure S1.** Deep learning pipeline for improving manually-generated brain masks.

**Table S1.** Cuttlefish brain lobe volumes.

