## Supplementary material for "A brain atlas of the camouflaging dwarf cuttlefish, *Sepia bandensis*": Figure S1 and Table S1

| Segment | Brain lobe | Number of voxels | Volume (mm3) | % of brain | |
|---|---|---|---|---|---|
| **AAB** | Anterior anterior basal lobe | 19314 | 2.41425 | | **0.77%** |
| **ADCl** | Anterior dorsal chromatophore lobe-L | 78 | 0.00975 | 0.00% | **0.01%** |
| **ADCr** | Anterior dorsal chromatophore lobe-R | 81 | 0.010125 | 0.00% | |
| **AP** | Anterior pedal lobe | 40469 | 5.05863 | | **1.62%** |
| **APB** | Anterior posterior basal lobe | 12074 | 1.50925 | | **0.48%** |
| **AVCl** | Anterior ventral chromatophone lobe-L | 227 | 0.028375 | 0.01% | **0.02%** |
| **AVCr** | Anterior ventral chromatophore lobe-R | 212 | 0.0265 | 0.01% | |
| **B** | Brachial lobe | 43001 | 5.37513 | | **1.72%** |
| **DB** | Dorsal basal lobe | 32291 | 4.03638 | | **1.29%** |
| **DLl** | Dorsolateral lobe-L | 1007 | 0.125875 | 0.04% | **0.08%** |
| **DLr** | Dorsolateral lobe-R | 1012 | 0.1265 | 0.04% | |
| **DMl** | Dorsal magnocellular lobe-L | 11896 | 1.487 | 0.48% | **0.95%** |
| **DMr** | Dorsal magnocellular lobe-R | 11901 | 1.48763 | 0.48% | |
| **DV** | Dorsal Vasomotor | 4757 | 0.594625 | | **0.19%** |
| **Fl** | Fin lobe-L | 10842 | 1.35525 | 0.43% | **0.87%** |
| **Fr** | Fin lobe-R | 10793 | 1.34913 | 0.43% | |
| **IBl** | Interbasal lobe-L | 1085 | 0.135625 | 0.04% | **0.09%** |
| **IBr** | Interbasal lobe-R | 1078 | 0.13475 | 0.04% | |
| **IF** | Inferior frontal lobe | 5114 | 0.63925 | | **0.20%** |
| **LBl** | Lateral Basal-L | 4504 | 0.563 | 0.18% | **0.36%** |
| **LBr** | Lateral Basal-R | 4511 | 0.563875 | 0.18% | |
| **LPl** | Lateral pedal lobe-L | 6198 | 0.77475 | 0.25% | **0.50%** |
| **LPr** | Lateral pedal lobe-R | 6175 | 0.771875 | 0.25% | |
| **LPVl** | Lateral ventral palliovisceral lobe-L | 4504 | 0.563 | 0.18% | **0.36%** |
| **LPVr** | Lateral ventral palliovisceral lobe-R | 4532 | 0.5665 | 0.18% | |
| **MB** | Median basal lobe | 32417 | 4.05213 | | **1.30%** |
| **OGl** | Optic Gland-L | 514 | 0.06425 | 0.02% | **0.04%** |
| **OGr** | Optic Gland-R | 528 | 0.066 | 0.02% | |
| **Ol** | Optic lobe-L | 939517 | 117.44 | 37.63% | **75.29%** |
| **Or** | Optic lobe-R | 940135 | 117.517 | 37.65% | |
| **OLl** | Olfactory lobe-L | 114 | 0.01425 | 0.00% | **0.01%** |
| **OLr** | Olfactory lobe-R | 120 | 0.015 | 0.00% | |
| **PC** | Precommisural lobe | 10720 | 1.34 | | **0.43%** |
| **PCLl** | Posterior chromatophore lobe-L | 3159 | 0.394875 | 0.13% | **0.25%** |
| **PCLr** | Posterior chromatophore lobe-R | 3128 | 0.391 | 0.13% | |
| **PF** | Posterior frontal lobe | 4495 | 0.561875 | | **0.18%** |
| **Pl** | Peduncle lobe-L | 7385 | 0.923125 | 0.30% | **0.82%** |
| **Pr** | Peduncle lobe-R | 7374 | 0.92175 | 0.30% | |
| **PMl** | Posterior magnocellular lobe-L | 5761 | 0.720125 | 0.23% | **0.46%** |
| **PMr** | Posterior magnocellular lobe-R | 5790 | 0.72375 | 0.23% | |
| **PP** | Posterior pedal lobe | 30019 | 3.75238 | | **1.20%** |
| **PV** | Palliovisceral lobe | 36475 | 4.55938 | | **1.46%** |
| **SF** | Superior frontal lobe | 25064 | 3.133 | | **1.00%** |
| **SV** | Subvertical lobe | 35773 | 4.47163 | | **1.43%** |
| **V** | Vertical lobe | 116763 | 14.5954 | | **4.68%** |
| **VMl** | Ventral magnocellular lobe-L | 2179 | 0.272375 | 0.09% | **0.18%** |
| **VMr** | Ventral magnocellular lobe-R | 2202 | 0.27525 | 0.09% | |
| **VV** | Ventral Vasomotor | 5721 | 0.715125 | | **0.23%** |

| Segment | Nerve tract | Number of voxels | Volume (mm3) | % of brain | |
|---|---|---|---|---|---|
| **AMC** | Anterior Magnocellular Commissure | 1237 | 0.154625 | | **0.05%** |
| **BPCl** | Brachio-pallioviscersal Connective-L | 2845 | 0.355625 | 0.11% | **0.23%** |
| **BPCr** | Brachio-pallioviscersal Connective-R | 2845 | 0.355625 | 0.11% | |
| **COl** | Collar Nerve-L | 111 | 0.013875 | 0.00% | **0.01%** |
| **COr** | Collar Nerve-R | 108 | 0.0135 | 0.00% | |
| **HRl** | Head Retractor Nerve-L | 677 | 0.084625 | 0.03% | **0.05%** |
| **HRr** | Head Retractor Nerve-R | 670 | 0.08375 | 0.03% | |
| **LBPCl** | Lateral Basal to Posterior Chromatophore Tract | 119 | 0.014875 | 0.00% | **0.01%** |
| **LBPCr** | Lateral Basal to Posterior Chromatophore Tract | 108 | 0.0135 | 0.00% | |
| **OABl** | Optic to Anterior Basal-L | 3641 | 0.455125 | 0.15% | **0.29%** |
| **OABr** | Optic to Anterior Basal-R | 3604 | 0.4505 | 0.14% | |
| **ODMl** | Optic to Dorsal Magnocellular-L | 1400 | 0.175 | 0.06% | **0.11%** |
| **ODMr** | Optic to Dorsal Magnocellular-R | 1409 | 0.176125 | 0.06% | |
| **OVl** | Optic to Vertical-L | 7419 | 0.927375 | 0.30% | **0.59%** |
| **OVr** | Optic to Vertical-R | 7400 | 0.925 | 0.30% | |
| **PNl** | Pallial Nerve-L | 2438 | 0.30475 | 0.10% | **0.19%** |
| **PNr** | Pallial Nerve-R | 2341 | 0.292625 | 0.09% | |
| **SOTl** | Subvertical to Optic Tract-L | 608 | 0.076 | 0.02% | **0.05%** |
| **SOTr** | Subvertical to Optic Tract-R | 603 | 0.075375 | 0.02% | |
| **VOC** | Ventral Optic Commissure | 4043 | 0.505375 | | **0.16%** |
| **VSl** | Visceral Nerve-L | 67 | 0.008375 | 0.00% | **0.01%** |
| **VSr** | Visceral Nerve-R | 65 | 0.008125 | 0.00% | |
| **SUM** | | | **312.096445** | | 100.00% |

**Table S1. Cuttlefish brain lobe volumes**

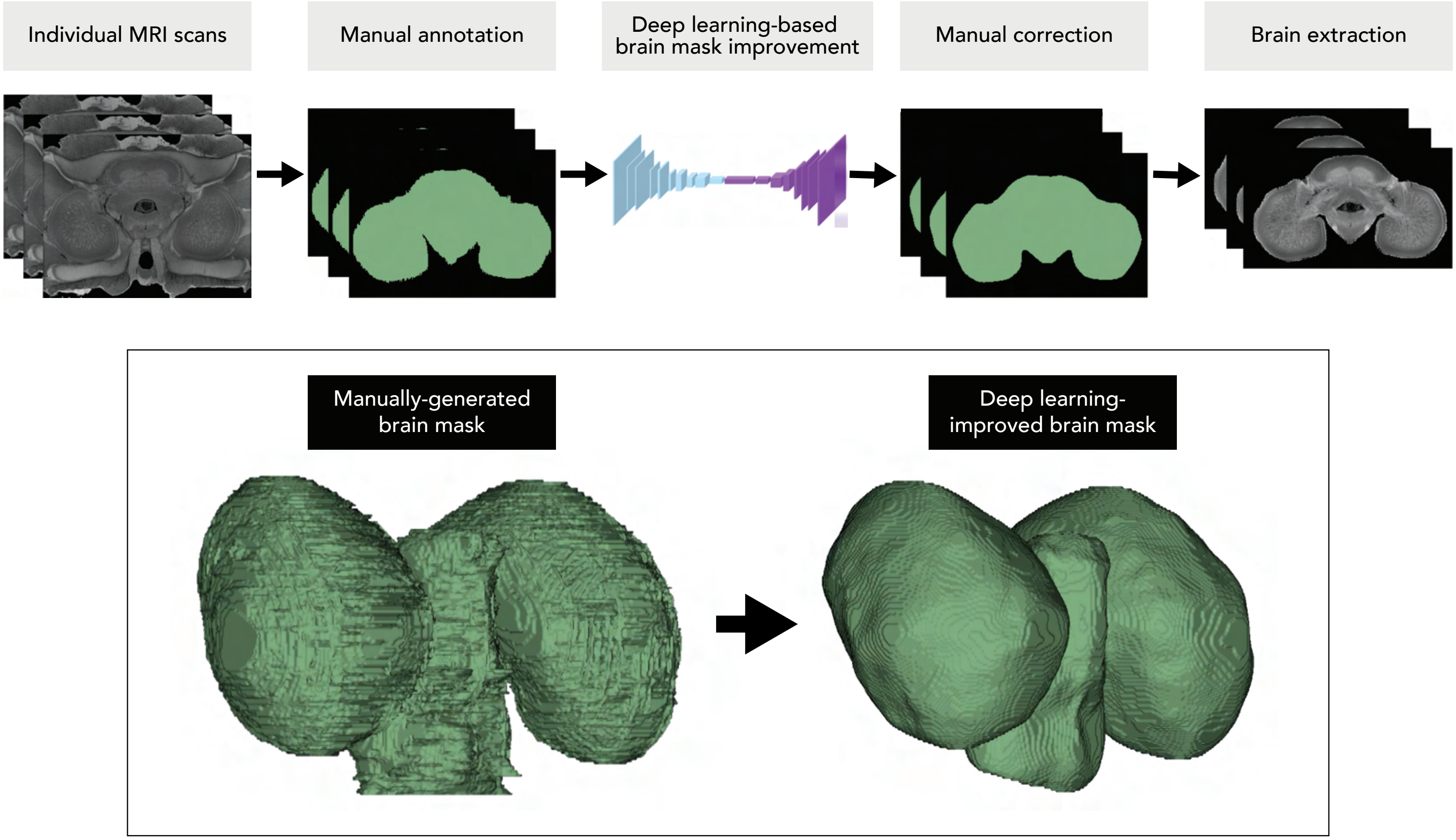


**Figure S1. Deep learning pipeline for improving manually-generated brain masks.**
